# Cortical travelling waves may underpin variation in personality traits

**DOI:** 10.1101/2025.01.15.633292

**Authors:** Neil W Bailey, Luiza Bonfim Pacheco, Luke D. Smillie

## Abstract

**Objectives:** Personality traits must relate to stable neural processes, yet few robust neural correlates of personality have been discovered. Recent methodological advances enable measurement of cortical travelling waves, which likely underpin information flow between brain regions. Here, we explore whether cortical travelling waves relate to personality traits from the “Big Five” taxonomy.

**Method:** We assessed personality traits and recorded resting electroencephalography (EEG) from 300 participants. We computed travelling wave strength using a 3D fast Fourier transform and explored relationships between alpha travelling waves and personality traits.

**Results:** Trait Agreeableness and Openness/Intellect had significant relationships to travelling waves that passed multiple-comparison controls (*p*_FDR_ = 0.019, *p*_FDR_ = 0.036). Agreeableness related to interhemispheric waves travelling from the right hemisphere along central lines (rho = 0.263, p < 0.001, BF10 = 356.350). This relationship was unique to the compassion aspect (t = 3.719, p <0.001) rather than politeness aspect of Agreeableness (t = 0.897, p = 0.370). Openness/Intellect related to backwards travelling waves along midline electrodes (rho = 0.197, p < 0.001, BF10 = 13.800), which was confirmed for the Openness aspect (rho = 0.216, p < 0.001, BF10 = 26.444) but not the Intellect aspect (rho = 0.093, p = 0.109, BF10 = 0.344).

**Conclusions:** Greater cortical travelling wave strength from right temporal regions may partly underpin variation in trait compassion, and backwards travelling wave strength along midline electrodes may mark trait openness. Further research is needed to investigate the mechanistic role of travelling waves in personality traits and other individual differences.

---

Personality traits are propensities toward particular patterns of emotion, cognition and behaviours that differ between individuals and are relatively stable across time (Wilt & Revelle, 2015). The dominant framework for measuring personality traits is the “Big Five” model (John et al., 2008), which organises personality trait variance using five core dimensions. These five dimensions comprise *agreeableness,* where high scoring individuals are characterised by compassion, politeness and interpersonal warmth, *neuroticism*, where high scoring individuals are characterised by anxiety, depression and emotional volatility, *conscientiousness*, where high scoring individuals are characterised as organised, hard-working and goal-oriented, *extraversion*, where high scoring individuals are characterised by sociability, enthusiasm and assertiveness, and *openness/intellect*, where high scoring individuals are characterised as showing imagination, curiosity, and aesthetic sensitivity. These traits are relatively stable over time, partly heritable, have been described across many cultures, and are predictive of a range of life outcomes (Bleidorn et al., 2014; Schmitt et al., 2007; Soto, 2019; Specht et al., 2011).

Given the stability and heritability of personality traits, it is sensible to assume that they have some relation to stable neurophysiological patterns. Accordingly, personality neuroscience research has investigated relations between personality traits and neural activity in an attempt to understand the mechanisms underlying these personality traits (Allen & DeYoung, 2017; DeYoung et al., 2022). For instance, resting electroencephalography (EEG) can be used to characterise trait-like neural activity at rest (Jach et al., 2020). EEG studies focussed on personality traits have commonly examined power within different frequency bands, which are typically assumed to reflect the brain’s oscillatory activity (Jach et al., 2020), but may also reflect the influence of aperiodic activity (Pacheco et al., 2024). These neural oscillations are commonly divided into different frequency bands (delta: 1-4Hz, theta: 4-8Hz, alpha: 8-13Hz, beta: 14-25Hz, gamma: >25Hz), which have been related to different cognitive processes (Buzsaki, 2006). For example, evidence suggests that alpha activity is related to the top-down inhibition of brain regions that are not required to complete current task demands (Klimesch et al., 2007; Wang et al., 2020). Some studies have found that specific frequencies relate to certain personality traits, for example posterior alpha power has been shown to negatively correlate with agreeableness (Jach et al., 2020), although a subsequent study failed to replicate this finding (Fruehlinger et al., 2024).

Typical analyses of EEG frequency power have measured only spatially stationary power, providing information only about voltage shifts assessed from a single point on the scalp. These measures might not sufficiently capture the multifaceted and complex neural activity related to personality traits (Fruehlinger et al., 2024). A growing area of research suggests that the voltage shifts that can be measured at a single electrode are in fact driven by cortical travelling waves, where voltage shifts progress across the cortex like waves traversing the ocean (Nunez, 1989; Visser et al., 2017). These travelling waves show amplitude peaks and troughs that travel through the cortex periodically (Alamia & VanRullen, 2019; Pang et al., 2020). Recent research indicates that cortical travelling alpha waves underpin information flow through the neural hierarchy, enabling attention and working memory functions (Bailey, Hill, Godfrey, Perera, Hohwy, et al., 2024; Lozano-Soldevilla & VanRullen, 2019; Mohan et al., 2024; Muller et al., 2014; Pang et al., 2020).

Furthermore, recent evidence suggests that cortical travelling alpha waves may be a mechanism by which the brain performs its “predictive processing” functions (Alamia et al., 2023). The predictive processing theoretical framework proposes that because our brains can only indirectly receive information about the environment (through sensory inputs), they function as a hierarchical Bayesian inference model generator to generate predictions about expected sensory inputs (Hohwy, 2020; Parr et al., 2022; Sprevak & Smith, 2023). Proponents of the predictive processing framework argue that non-predicted sensory inputs (prediction errors) are passed up the neural hierarchy to update our brain’s predictive model, while predicted inputs are suppressed (Hohwy, 2020; Parr et al., 2022; Sprevak & Smith, 2023). Active inference is then suggested as the mechanism by which the brain acts on its environment (via muscle control) to match the environment to its predictions, enabling goal directed behaviour (Parr et al., 2022). This parsimonious theory is rapidly becoming the dominant paradigm in neuroscience for its elegant explanation that can apply across behaviours and brain functions.

The potential explanatory power of predictive processing and cortical travelling waves might be enhanced by our understanding of the function of different brain regions. Sensory inputs are predominantly processed in posterior brain regions lower in the cortical hierarchy (Aitken et al., 2020; Gordon et al., 2019), while frontal regions that are higher in the cortical hierarchy primarily generate more abstract predictions about expected sensations (Hodson et al., 2023; Sprevak & Smith, 2023). Top-down processes higher in the cortical hierarchy are suggested to relate to more complex and abstract predictive modelling, including thoughts about past events and future events, whereas bottom-up processes reflect more sensory processing (Corcoran et al., 2020; Laukkonen & Slagter, 2021). Individual differences in the balance of this functional dichotomy might relate to personality traits such as trait openness/intellect, which has known associations to executive functions like voluntary attention control (DeYoung et al., 2014), a function that may be implemented via stronger application of top-down prediction weightings within this predictive processing framework (Alamia & VanRullen, 2019; Pang et al., 2020). Thus, Openness/Intellect might be expected to relate to stronger backwards travelling waves, given that individuals higher on this trait tend to exhibit higher cognitive ability, working memory, creative problem solving, and engagement of attention and executive functions (DeYoung et al., 2014; Murdock et al., 2013; Smillie et al., 2016). Similarly, inferences about the functional relevance of specific brain regions combined with measures of information flow from those regions might provide utility in understanding the mechanisms underlying personality traits. For example, activity in the right temporoparietal junction has been shown to be associated with the processing of empathy, sympathy, perspective taking, and social cognition (Decety & Lamm, 2007). As these processes are relevant to trait agreeableness, it seems plausible that individual variation in information flow from the right temporoparietal junction could be a driver of differences in this trait. Thus, Agreeableness might correlate with travelling waves from regions that relate to processing the emotions and intentions of others, for example the temporoparietal junction (Gallagher et al., 2000). However, to our knowledge, no research has explored how cortical travelling wave patterns may relate to any personality traits.

Regarding the measurement of travelling waves, oscillations in posterior electrodes that are also found in more anterior electrodes, but with phase delays progressively increasing as the electrodes become more anterior, are assumed to be waves travelling forwards (Alamia & VanRullen, 2024). In contrast, phase lags that progressively increase from frontal to posterior electrodes suggest a backwards travelling wave (Alamia & VanRullen, 2024). Backwards travelling waves have been shown to reflect top-down predictions and to reflect the direction of voluntary attention (Alamia & VanRullen, 2019; Pang et al., 2020). Forwards travelling waves have been shown to relate to the bottom-up propagation of prediction errors related to sensory inputs (Alamia & VanRullen, 2019; Ermentrout & Kleinfeld, 2001; Pang et al., 2020). Thus, cortical travelling waves capture information flow through the cortex and provide information about the explanatory predictive processing functions of the brain, providing a novel avenue for personality neuroscientists investigating biological mechanisms underlying personality traits.

We therefore undertook an exploratory analysis of a large existing dataset, measuring alpha waves travelling both forwards and backwards and laterally across the scalp, and relating these to the Big Five personality traits. This may help identify potential neural mechanisms underpinning personality, and potentially offer greater interpretability than previously identified associations between neural activity and personality traits.

## Methods

### Participants and Procedure

The dataset comprised a total of 300 right-handed participants aged between 18 and 60 years, who were recruited to multiple studies through physical and online notices. The full sample has been described by Pacheco et al. (2024) and a subset of the data was described earlier by Jach et al. (2020). All participants had normal or corrected-to-normal vision and did not report any current mental illness.

After pre-processing the EEG data, 296 participants (176 female, 120 male) provided enough artifact free electrodes and epochs for inclusion in the final analysis (four participants were excluded due to an excessive number of bad electrodes, with the details of these exclusion criteria provided in the Procedure section). The mean age of the participants for the final sample was 23.108 (SD = 6.275).

Ethical approval for the study was provided by the Human Research Ethics Committee of the University of Melbourne (ID 1954069). All participants provided written informed consent prior to participation in the study.

Prior to the EEG recording and task completion, participants provided demographic information and completed the self-report personality questionnaire described below via an online *Qualtrics*^TM^ survey. Demographics included age, sex and hand preference.

### Personality Assessment

All participants completed the Big Five Aspects Scales (DeYoung, Quilty and Peterson, 2007), which comprises 100 items to assess the Big Five at two levels of abstraction: The *domain* level comprises openness/intellect, neuroticism, agreeableness, conscientiousness, and extraversion (20 items each). Each of these domains divides into two *aspect* level traits; the ten aspects are openness, intellect, volatility, withdrawal, politeness, compassion, orderliness, industriousness, assertiveness, and enthusiasm (10 items each). Each item consists of a brief descriptive phrase (e.g., for extraversion: *make friends easily*) to which participants respond by indicating their level of agreement that the phrase describes their personality (from 1 = *Strongly Disagree* to 5 = *Strongly Agree*).

### EEG Acquisition and Pre-Processing

Sixty-four-channel EEG (BioSemi, Amsterdam, The Netherlands) was recorded from Ag/AgCl electrodes embedded in an elasticized Easy-Cap® and aligned with the extended 10–20 system. Additional electrodes were also placed at the outer canthus and supra-orbit of the left eye, and at the right and left mastoids, although these electrodes were not used in the current analysis. All electrodes’ offsets were within ±40µV, and EEG data were sampled at 512Hz with no online bandpass filters. Participants rested with 8 interleaved periods of 60s with either their eyes open or eyes closed; the experimenter instructed participants to switch between eyes open and eyes closed when a visual message signalled the end of each 60s window. Electrodes were grounded using common mode sense and driven right leg electrodes.

EEG data were pre-processed using the RELAX EEG pre-processing pipeline (Bailey et al., 2023b), a toolbox that uses EEGLAB functions (Delorme & Makeig, 2004), with selected application of Fieldtrip functions (Oostenveld et al., 2011). The RELAX cleaning pipeline first bandpass filters the data between 0.25Hz and 80Hz, with a notch filter from 47 to 53Hz, using fourth order Butterworth filters. The pipeline then uses the PREP toolbox to reject bad electrodes (Bigdely-Shamlo et al., 2015), followed by a secondary rejection of electrodes that show outlying data for more than 5% of the recording using moderately aggressive outlier rejection settings, with outliers detected using measures of kurtosis, the probability of the distribution of voltage values within each epoch, absolute voltage thresholds and voltage shifts within an epoch (Bailey, Hill, Godfrey, Perera, Rogasch, et al., 2024), with the limit that no more than 10% of electrodes could be rejected. Outlying periods remaining in the data were then rejected based on the same outlier identification approaches as the channel rejection step.

Following the rejection of bad electrodes, three sequential multi-channel Wiener (MWF) filters were applied as an initial artifact rejection step to reduce muscle artifact, then blinks, then both horizontal eye movement and remaining drift. To clean EEG data with MWF, the first step involves obtaining templates of EEG data periods that are identified as containing artifacts and periods identified as containing only neural activity. MWF then uses a delay embedded matrix of data from all electrodes in combination with the artifact and clean data templates to obtain a model of the spatio-temporal patterns that characterize the artifacts through a minimization of the mean squared error algorithm (Somers et al., 2018). Here, we used a delay embedding period set to 16, which means that the MWF cleaning characterised each artifact by accounting for a total of 33ms of data at a time (16 samples forwards and backwards from each timepoint). Since the delay embedding characterizes the temporal as well as spatial aspects of the artifacts, MWF acts as a spatio-temporal filter. This allows MWF to reduce artifact topographies by taking into account the temporal patterns as well as the spatial patterns of the artifacts (Somers et al., 2018). This has been suggested to effectively clean artifacts while also better preserving neural activity (Somers et al., 2018).

Following the MWF cleaning, PREP’s robust average referencing was applied to the data (Bigdely-Shamlo et al., 2015). Independent component analysis (ICA) was then computed on the data using the fastica algorithm. Artifactual components remaining in the data were identified by the machine learning algorithm, ICLabel, with the criteria that if ICLabel identified a component as “most likely” to be artifact, it was selected for reduction (Pion-Tonachini et al., 2019). Artifacts from all categories were identified (eye movements/blinks, muscle activity, heart-beat, line noise, and non-specific other artifacts). These artifacts were then reduced using a stationary wavelet transform to characterize the dominant frequencies within each artifact component, which are assumed to reflect the artifact contribution to the signal, and removed from each component before data were reconstructed back into the scalp space (Castellanos & Makarov, 2006). Missing electrodes were interpolated back into the data using spherical spline interpolation (Perrin et al., 1989). This combination of MWF and wavelet enhanced ICA has been shown to effectively clean EEG data of artifacts, while preserving neural signals (Bailey et al., 2023b; Bailey et al., 2023c).

Following cleaning of the continuous data, data were epoched into one second periods with a 0.5s overlap from the eyes open periods of the recording. Selected electrodes from these epochs were then extracted to enable computation of the cortical travelling waves (electrodes and computation method described in detail below).

### Measures

Personality scores were computed by taking the mean response to the 20 items for each domain and the 10 items for each aspect, for a total of 15 scale scores. We also estimated these scores as latent variables (using maximum likelihood estimation), which reduces measurement error (because only the variance shared by all items within a respective scale is represented). As results based on latent variables did not differ substantively from those based on mean scores we relegate theses analyses to the supplement.

To determine the strength and direction of cortical travelling alpha travelling waves, we expanded the method developed by Alamia and VanRullen (2019) to include 35 electrodes across both lateral and midline locations from both hemisphere (AF7, AF3, AFz, AF4, and AF8, as well as PO7, PO3, POz, PO4, and PO8, and all electrodes between). This allowed the measurement of cortical travelling waves in both forwards and backwards and lateral directions. See Figure 1 for a visual depiction of this methodology. Within each participant, 1 second epochs were extracted from each of the EEG recordings. Following the methods introduced by Alamia and VanRullen (2019), these epochs were extracted every 500ms within the resting data (so the epochs from the resting data contained 500ms overlaps with neighbouring epochs). This provided a 3-dimensional matrix for each epoch (forwards and backwards electrode line x lateral electrode line x time). A 3-dimensional fast Fourier transform (3D-FFT) was then applied (using the matlab function ‘fftn’) to the 3D matrix from each epoch separately. The output matrix of this 3D-FFT provides both temporal frequencies and spatial frequencies, with spatial frequencies represented in the first two axes of the matrix (see Figure 1). Waves propagating in the forwards direction (from occipital to frontal electrodes) are represented in the upper quadrant of the matrix (Figure 1C), while waves propagating in the backwards (from frontal to occipital electrodes) are represented in the lower quadrant (Alamia & VanRullen, 2019). Within the 3D version of the matrix, waves propagating from right to left are represented in the left side of the matrix, and waves propagating from left to right are represented in the right side of the matrix (the 3D version of the matrix is not depicted in Figure 1 due to its complexity).

**Figure 1.**
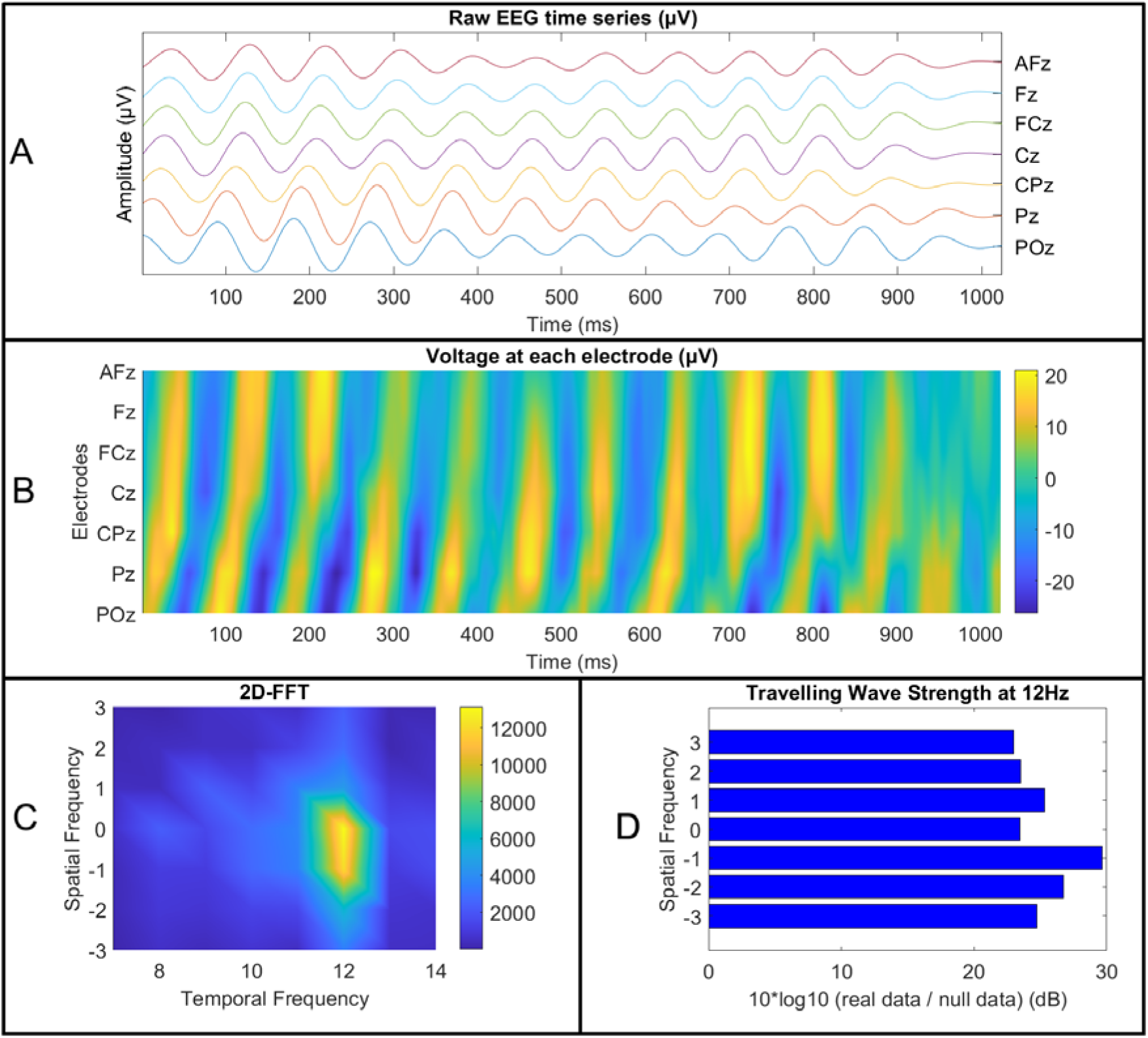
A visual representation of the travelling wave computation method using a 2-dimensional fast Fourier transform (2D-FFT), which can be extrapolated to understand the 3-dimensional fast Fourier transform (3D-FFT) used in our primary analyses. First, 1 second epochs were extracted from the continuous electroencephalography (EEG) data after removal of artifacts. The time-series from each electrode were then organised into matrices, with posterior electrodes at the bottom of the matrices and anterior electrodes at the top (and left electrodes on the left side of the 3D matrices, and right electrodes on the right side), as depicted in A and B (after applying a bandpass filter in the alpha frequencies of interest to highlight the travelling waves). Each epoch was then used as the input for a 3D-FFT (MATLAB), or 2D-FFT in the case of our post-hoc tests (as depicted above for simplification of the visualisation). This provided a matrix providing power values in each spatial and temporal frequency. Within this matrix, the x-axis provides power values at each temporal frequency, while the y-axis provides power values at each spatial frequency – shown in C with values above 0 reflecting backwards travelling spatial waves (from frontal to posterior electrodes), values below 0 reflecting forwards travelling spatial waves (from posterior to frontal electrodes), and values at 0 reflecting standing waves (waves that do not travel). In our 3D-FFT (not depicted in this figure), the z-axis would reflect lateral travelling waves (from left to right or right to left). Finally, to assess the travelling wave strength in each spatial and temporal frequency, we divided the power in each cell in the matrix by the power obtained after randomly shuffling the order of the electrodes in each epoch 100 times, repeating the FFT computations for each shuffle, and averaging the outputs of these null FFTs. We then computed a log10 transform on the output of this division and multiplied this value by 10 to obtain a measure of the strength by which the real travelling wave strength exceeded the permutation-based null travelling wave strength, on the decibel scale (Figure 1D).

To ensure these travelling wave values reflected real signals that exceeded simple random patterns in the data, and to control for the potential influence of variation in oscillatory power without variation in travelling wave strength, we performed the same travelling wave computations on 100 surrogate null versions of the data for each epoch. These surrogate null versions were obtained by randomly shuffling the location of all electrodes in the matrix from each epoch separately prior to the computation of the 3D-FFT on this null data (Alamia & VanRullen, 2019). This shuffling of the electrode order made it impossible for the analysis to detect the spatial pattern of travelling waves provided by progressively increasing phase lags between electrodes, yet preserved the temporal oscillatory pattern, providing a distribution that was matched to the real data for oscillatory power but reflecting a null distribution for the spatial structure (Alamia & VanRullen, 2019). Then, to ensure our statistical comparisons focused on the signal strength of the real travelling waves above the null, we performed a matrix-wise division of the values from the real data within each epoch by the values within the surrogate data for the same epoch, then multiplied the result by 10*log10. This provided a value for each epoch within each participant that reflected the ratio of the strength by which the real travelling waves exceeded the surrogate waves, with values on a log scale (providing values in decibel units [dB]) (Alamia & VanRullen, 2019). Finally, we extracted the 75^th^ percentile travelling wave value within each cell in the matrix, inclusive of backwards and forwards travelling waves at three spatial frequencies, left and right travelling waves at two spatial frequencies, and six temporal frequencies (8 to 13Hz). The 75^th^ percentile value was used instead of the mean, as we reasoned that potential differences in travelling wave strength between different personality traits would likely be driven by the engagement of cortical travelling waves, reflecting active neural processing related to a personality trait, more than by the average across all resting neural activity. However, we note that the significant effects presented in our results section remained significant when using the mean value instead of the 75^th^ percentile, albeit with a reduced effect size (reported in our supplementary materials).

### Statistical Analyses

Analyses were performed using MATLAB (Mathworks, 2023a), R (R-Core-Team, 2024), and JASP 0.17.2.1(Love et al., 2019). The output of our 3D-FFT provided 35 spatial frequencies and 6 temporal frequencies, which could each be correlated against 5 personality traits (and 10 sub-traits). Given the large number of potential multiple statistical tests with likely dependence between different tests, we used cluster-based statistics, which do not assume independence between tests, but rather leverages the likely dependence between tests to increase the power to detect real effects against a null distribution obtained by shuffling individual labels. First, we performed Spearman’s correlations between each output from the 3D-FFT and each personality trait to obtain a single 3D matrix of rho values for each personality trait. Next, we excluded non-significant correlations (p > 0.05) from each 3D matrix of rho values. We then computed the maximum sum of rho values from three adjacent cells, across at least two adjacent frequencies and two adjacent spatial cells that passed our initial threshold for each of these 3D matrices (p < 0.05). Following this, we performed an absolute transform on the maximum from the 3D rho matrix for each personality trait. Next, we ranked these absolute maximum sum rho values from significant clusters against the absolute maximum sum rho values from clusters that passed our initial threshold obtained from 5000 shuffles of the participant labels. This enabled us to obtain an overall p-value for the relationship between each personality trait and the overall cortical travelling wave matrix. Clusters from the real data that provided larger absolute maximum sum values than 95% of the null distribution data were deemed significant at p < 0.05. Finally, we submitted the overall p-value for each personality trait to an experiment-wise false discovery rate (FDR) multiple statistical test control using the method introduced by Benjamini and Hochberg (1995).

Following our tests using cluster-based statistics to determine whether any personality traits correlated to any cortical travelling waves as measured by the output of our 3D-FFT computation, we performed post-hoc tests of relationships between the personality traits showing effects in our overall analysis using 2D-FFT restricted to the patterns that showed the strongest effects in our 3D-FFT. This enabled us to determine the specific pattern of cortical travelling waves that related to each personality trait (since cluster-based statistics can provide an overall measure of significance, but interpretation of the significance of the location of an effect using cluster-based statistics is not valid). Within these 2D-FFT analyses, we first obtained the maximum travelling wave strength within the temporal frequencies of interest (those showing a significant effect within the cluster analyses of the 3D-FFT), which enabled us to reduce the number of comparisons. Then we performed the same log10 normalisation as described for the 3D-FFT to obtain the 2D travelling wave strength for each individual in each effect of interest.

We then used Spearman’s correlations to examine relationships between personality scores and the specific travelling wave directions and frequencies that showed significant effects within our cluster analysis of the 3D-FFT, and Bayesian Pearson’s correlations between the same variables to obtain Bayes Factors (BF) enabling us to determine the strength of evidence for the null or alternative hypotheses (BF10 indicates the probability that the alternative hypothesis is accurate given the data compared to the probability that the null hypothesis is accurate given the data).

## Results

Domain-level Agreeableness and Openness/Intellect showed relationships to cortical travelling waves in our eyes open resting data that passed our cluster and Benjamini Hochberg FDR multiple comparison controls (*p*_FDR_ = 0.019 and *p*_FDR_ = 0.036 respectively, Figure 2). None of the other personality factors passed our statistical thresholds: Neuroticism *p*_FDR_ = 0.299, extraversion *p*_FDR_ = 0.064, and conscientiousness *p*_FDR_ = 0.463.

**Figure 2.**
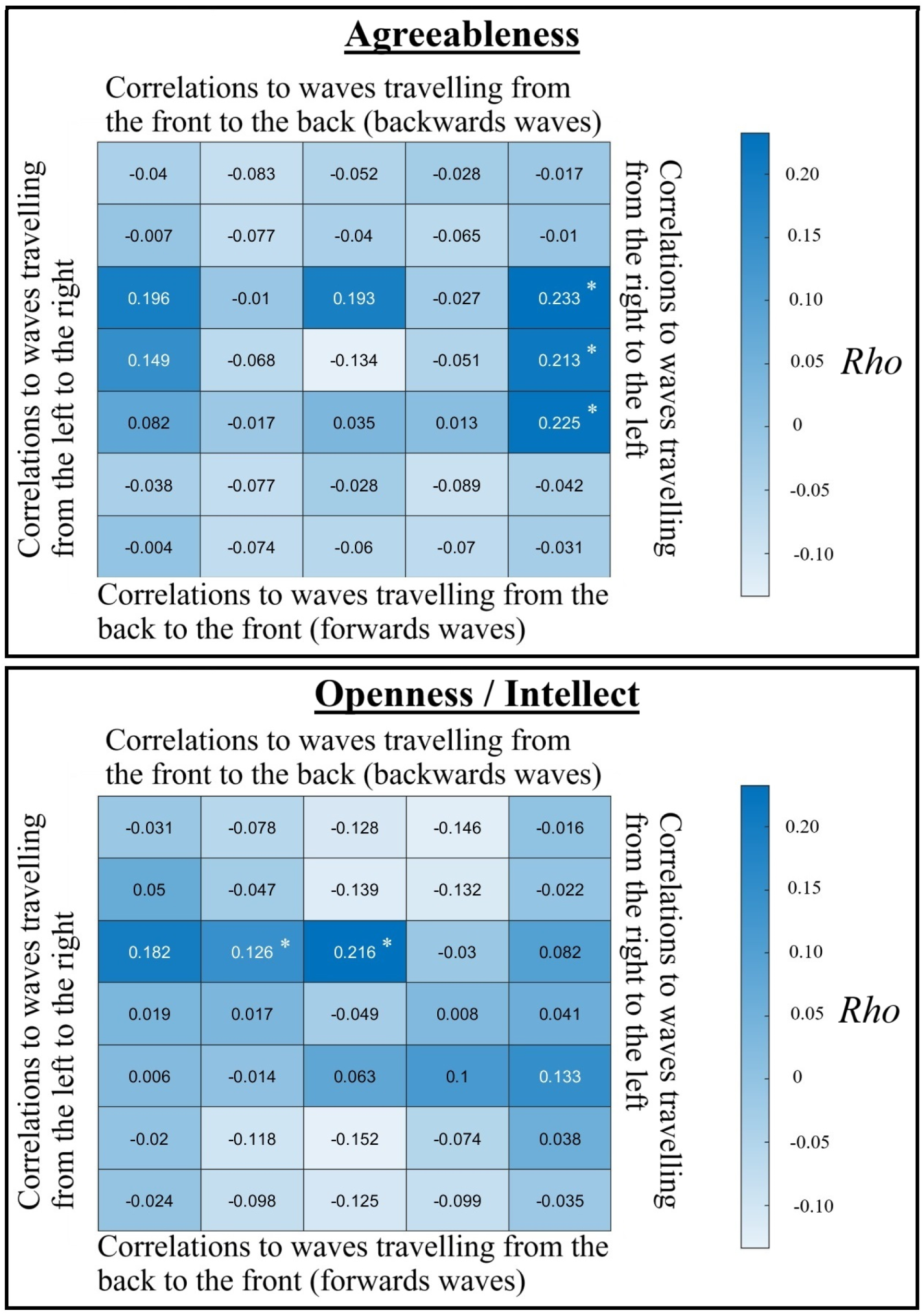
Spearman’s correlations between 3D-FFT travelling wave values at each spatial frequency and direction, and at 10Hz in the frequency domain, and agreeableness (top) or openness / intellect (bottom). * indicates cells that were involved within a cluster that passed our initial cluster statistical test of relationships between the personality traits and 3D-FFT outcomes (*p*_FDR_ < 0.05), with values in each cell representing travelling waves across the directions that could be measured across the scalp electrodes.

The cluster showing the strongest rho values for correlations between Agreeableness and cortical travelling waves in our 3D analysis were for waves travelling from the right to the left hemisphere at the highest spatial frequency, along the central, fronto-central, and centroparietal electrode lines, which showed a positive correlation with maximal rho values between 9-10 Hz in the temporal frequency dimension (Figure 2), but also significant effects from 8-13Hz within this cluster. After excluding the cluster showing the largest effect, an additional cluster from 8-9Hz travelling left to right at the maximal spatial frequency and from 8-11Hz in the temporal frequency dimension also passed our FDR threshold, showing a positive correlation with Agreeableness (*p*_FDR_ = 0.015).

The cluster showing the strongest rho values for correlations between Openness/Intellect and cortical travelling waves in our 3D analysis were for backwards travelling waves at the lowest spatial frequency along the midline electrodes with maximal rho values from 10-11Hz, and significant effects from 9-12Hz. These clusters showed positive correlations with Openness/Intellect (Figure 2).

### Agreeableness Post-hoc Tests

Exploration of the significant effects within the cluster statistics used to examine the 3D-FFT outcomes with post-hoc 2D tests showed that Agreeableness significantly correlated to waves travelling from the right to the left hemisphere along the central electrode lines (T8, C4, Cz, C4, T7) in the 9-10Hz temporal frequencies (*rho* = 0.263, *p* < 0.001, *BF10* = 356.350, Figure 3). To test the potential replicability of this result, we randomly split the data into half the sample size 10 times, then tested Spearman’s correlations on each of these reduced samples. All of the ten reduced samples showed a significant correlation (all *p* < 0.005, mean *p* < 0.001, all *rho* fell between 0.237 and 0.344, mean *rho* = 0.299). Exploring potential relationships between this rightwards travelling wave and the different aspects of agreeableness, we found that the association was appreciably stronger for the compassion aspect of Agreeableness (*rho* = 0.285, *p* < 0.001, *BF10* = 810.438, Figure 3) than the politeness aspect (*rho* = 0.167, *p* = 0.004, *BF10* = 1.437, Figure 3), although Hittner et al.’s (2003) test for the difference between dependent correlations with overlapping variables fell just short of formal significance (Z = 1.912, *p* = 0.056). Given the overlap between compassion and politeness (*rho* = 0.401, *p* < 0.001), a linear regression including compassion and politeness as predictors for the rightwards cortical travelling wave strengths was undertaken. This yielded a significant effect for compassion (*t* = 3.719, *p* < 0.001) but not for politeness (*t* = 0.897, *p* = 0.370), suggesting the effect was specific to compassion.

**Figure 3.**
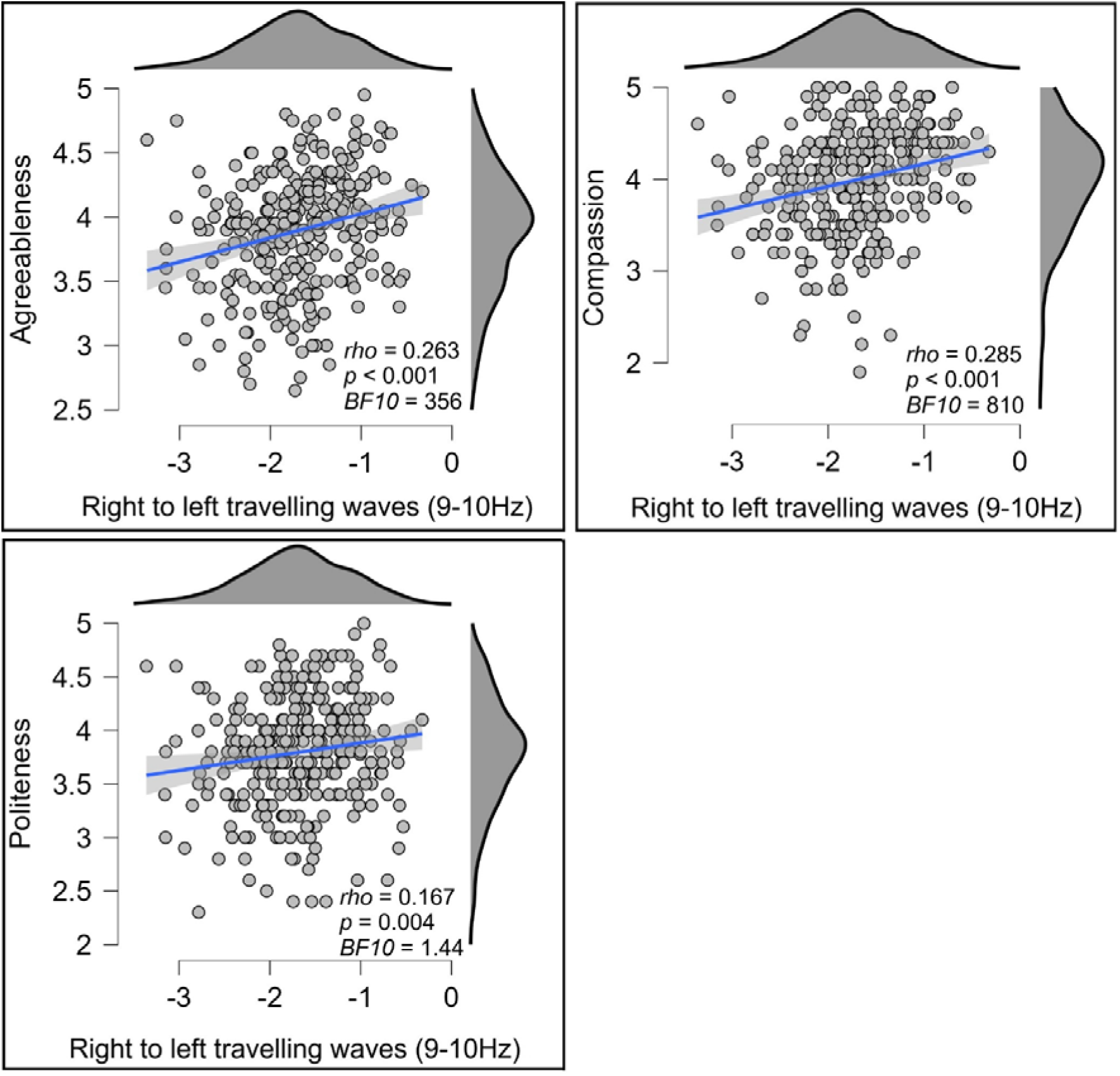
Scatterplots showing the relationship between agreeableness (top left), compassion (top right), or politeness (bottom left) and right to left cortical travelling alpha waves (at 9 to 10Hz, computed using a 2D-FFT from the central electrode line, units in decibels) within the eyes open resting data. The blue lines indicate the linear regression line with the shaded area indicating the 95% confidence interval for this regression line.

In addition to the significant rightwards travelling wave cluster in our cluster-based statistical analysis performed the 3D-FFT outputs, a cluster from 8-9Hz travelling left to right at the maximal spatial frequency showed a similar positive correlation with Agreeableness. This cluster also showed a similar positive correlation with Agreeableness when examined in the 2D analysis along the central electrodes, although with Bayesian evidence against the alternative hypothesis (*rho* = 0.155, *p* = 0.007, *BF10* = 0.718). The significant effect was also present for compassion (*rho* = 0.158, *p* = 0.006, *BF10* = 0.608), but not politeness (*rho* = 0.106, *p* = 0.068, *BF10* = 0.215); the difference between these effects was not significant, Z = 0.823, *p* = 0.411. Further, a linear regression including compassion and politeness as predictors for the leftwards travelling wave strengths did not yield a significant effect for either compassion (*t* = 1.629, *p* = 0.104) or politeness (*t* = 0.744, *p* = 0.458).

Next, to assess the specificity of the rightwards travelling wave effects, and their robustness to different analytical choices, we performed several validation analyses. For brevity, the full report of these results is restricted to our supplementary materials, and we only provide a summary here. First, testing the same relationships to rightwards travelling waves during eyes closed resting data (rather than eyes open resting data) showed the same pattern of results and same significant effects, indicating that the patterns are consistent across different resting states. Second, when we tested correlations between the rightward travelling wave and latent variables instead of mean responses, the exact same pattern of results and significant effects emerged. We also tested relationships between cortical travelling waves along the fronto-central, and centroparietal lateral electrode lines and agreeableness (and its aspects). These results showed the exact same pattern of results and significant effects for the centroparietal electrode line (TP7 to TP8), indicating a broad effect from the electrodes over centro/temporo-parietal regions. The analysis of the fronto-central line (FT7 to FT8) showed a similar pattern of results, but with weaker effect sizes. Additionally, the results of the regression analysis suggested that neither compassion nor politeness was predictive of rightwards travelling wave strength (although there was a non-significant trend effect for compassion in the same direction as our results for the other electrode lines, *t* = 1.919, *p* = 0.056).

Next, we noted that values for the strongest spatial frequency in the rightwards lateral travelling waves were negative after performing the log transform on the real values divided by the mean of 100 null shuffles. This suggests that the null shuffles of electrode orders contained more power than the real data in these highest lateral spatial frequencies. This pattern might be explained by an increased strength of travelling waves between the midline electrodes compared to waves propagating from outer lateral electrodes, such that when the midline electrodes are “spread out” in the null shuffles of the data, the increased strength of travelling waves between midline electrodes is transferred to the higher spatial frequencies. As a result, the log transform of the real values divided by the null shuffle values produces negative values on the decibel scale in the normalised data. This prompted us to assess whether the positive correlations we detected were driven by the presence of true lateral travelling waves rather than somehow being driven by the normalisation procedure.

Results of these assessments showed that the relationships to agreeableness were indeed driven by variability in actual rightwards travelling wave strength. Positive and significant correlations with the same patterns as our primary analyses were still obtained in tests of data that were normalised against mean alpha power across the lateral electrodes, obtained using a 1D FFT. This approach controls for variations in alpha power without being influenced by patterns in the null shuffle data (Zeng et al., 2024). Positive and significant correlations with the same patterns as our primary analyses were also obtained in tests of the data normalised against nulls obtained by shuffling electrodes across epochs (rather than shuffling the order of electrodes within the epoch). This approach preserves variations in relative alpha power between electrodes in the null data, and controls for overall alpha power, so this approach restricts tests to the variability that is specific to the spatiotemporal patterns (phase and power relationships between electrodes) that indicate real travelling waves in the real data. Interestingly, normalising using this method showed rightward travelling waves exceeded the null shuffled data in many participants, but not all participants, with a mean value of 0.170 dB (*SD* = 0.306, minimum = -0.786, maximum = 0.885).

Given the results in these validation tests, we can conclude that, although the rightwards travelling wave reflects a true signal in the data for many (but not all) individuals, the signal is weak. These tests confirm that the effects were driven by a relationship between true (but weak) rightwards travelling waves. These rightwards travelling waves were present more commonly in individuals who scored higher in agreeableness, and individuals who scored higher in agreeableness showed rightwards travelling waves that exceeded the values obtained via null shuffles by a larger amount compared to individuals who scored lower in agreeableness, particularly for the compassion aspect. In contrast, the mean of the null shuffles of the data did not correlate with agreeableness or its compassion or politeness aspects, confirming that the normalisation against the null shuffle data was not driving the detected effects. These results are reported in full in the supplementary materials.

Furthermore, since previous research on a subset of the dataset used in the current study found that agreeableness is negatively correlated with posterior alpha power (Jach et al., 2020), we examined whether our results might be simply driven by differences in alpha power. To test this, we z-score transformed each electrode’s time series separately prior to the 2D-FFT. This normalises for differences in amplitude between electrodes and between individuals, controlling for potential differences in alpha power, while preserving the spatial properties of the cortical travelling waves (since the phase angles between the electrodes are preserved by this transform). The full results of this analysis are also reported in our supplementary materials; in brief, the same pattern of results and significant effects was present as per our primary analyses reported above, suggesting the effects were driven by variation in travelling wave strength, rather than simply a relationship between alpha power and agreeableness.

We note that the analyses of lateral travelling waves reported thus far do not reveal whether the waves travel within the right hemisphere (from the temporal region towards the midline), or whether they travel between hemispheres. To address this, we performed additional 2D-FFTs of selected electrodes, including only right hemisphere electrodes, only electrodes closer to the midline, or the full set of 9 potential central electrodes. Relationships between Agreeableness and lateral cortical travelling waves restricted to the right hemisphere only were not significant (all *p* > 0.10), suggesting our primary results were not driven by waves travelling only within the right hemisphere. Similarly, when our 2D-FFT was restricted to the more midline electrodes (from C3 to C4), no significant correlations were present (all *p* > 0.10). Only when we included electrodes from C5 to C6 or when we included all 9 central line electrodes did significant effects become apparent, with a significant correlation between Agreeableness and right to left travelling waves similar to our primary results (*rho* > 0.164, *p* < 0.005, *BF10* > 3.800). This pattern of results suggests that the relationship between right to left cortical travelling waves and agreeableness / compassion is produced by interhemispheric cortical travelling wave patterns rather than travelling waves from lateral electrodes to midline electrodes. Finally, our exploratory analysis of the data quantified using the mean value across epochs (rather than the 75^th^ percentile) showed the same patterns and significant results as our primary analyses (reported in full in the supplementary materials).

### Openness/Intellect Post-hoc Tests

Our 2D-FFT restricted to the midline electrodes (POz, Pz, CPz, Cz, FCz, Fz, AFz) showed a significant positive correlation between Openness/Intellect and backwards travelling waves at the lowest spatial frequency and 10-11Hz in the temporal frequency (*rho* = 0.197, *p* < 0.001, *BF10* = 13.800, Figure 4). To test the potential replicability of this result, we randomly split participants into separate groups of half the total sample size 10 separate times, then tested Spearman’s correlations on each of these reduced samples. Nine of the ten reduced samples showed a significant correlation (all *p* < 0.054, 9/10 *p* < 0.030, mean *p* = 0.022, all *rho* fell between 0.159 and 0.274, mean *rho* = 0.197). Given the weaker correlation strength than for relationships to Agreeableness, we suspected the fact that significant effects were present for only 9/10 splits might be due to a smaller sample size in the half sample splits. To assess this, we instead tested whether the correlations would remain significant if we used 75% of the sample in our splits. All of these ten reduced samples showed a significant correlation (all *p* < 0.036, mean *p* = 0.009, all *rho* fell between 0.141 and 0.224, mean *rho* = 0.194).

**Figure 4.**
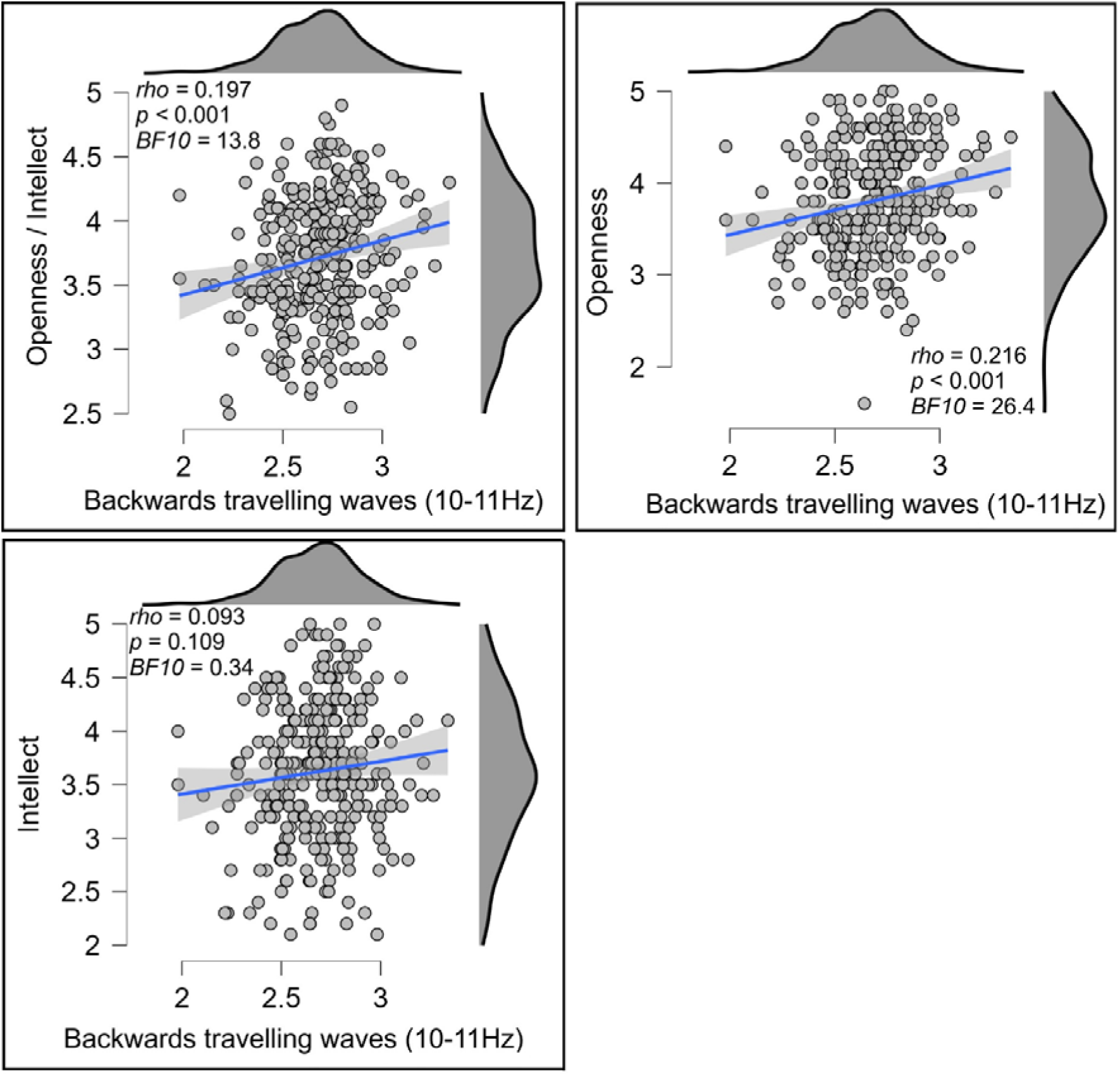
Scatterplots showing the relationship between Openness / Intellect (top left), openness (top right), or intellect (bottom left) and backwards travelling alpha waves (at 9 to 10Hz, computed using a 2D-FFT from the midline electrodes, units in decibels) within the eyes open resting data. The blue lines indicate the linear regression line with the shaded area indicating the 95% confidence interval for this regression line.

In exploring whether the effect was related to specific aspects of Openness/Intellect, we found that the correlation was stronger for the Openness aspect (*rho* = 0.216, *p* < 0.001, *BF10* = 26.444, Figure 8), and that the correlation with the Intellect aspect was not significant (*rho* = 0.093, *p* = 0.109, *BF10* = 0.344, Figure 9); the difference between these correlations fell short of formal significance, Z = 1.768, *p* = 0.077. Exploring further, a linear regression including these two variables as predictors for the backwards cortical travelling wave strengths revealed a significant effect for openness (*t* = 3.128, *p* = 0.002) but not for intellect (*t* = 0.956, *p* = 0.340), suggesting the effect was unique to openness.

Furthermore, in addition to the relationship between backwards travelling waves and Openness/Intellect in the eyes open data, we tested for relationships to the eyes closed resting data to determine the consistency of the effect across different states. The same correlations between travelling waves and Openness/Intellect and the Openness aspect were significant in the eyes closed data, although with only weak Bayesian evidence in support of the relationship to Openness/Intellect (reported in full in the supplementary materials). These results indicate that the patterns were consistent across different resting states. Finally, our exploratory analysis of the data quantified using the mean value across epochs (rather than the 75^th^ percentile) showed the same patterns and significant results as our primary analyses (reported in full in the supplementary materials).

## Discussion

In this study we used a novel 3D-FFT analysis to determine whether any personality traits were related to the strength of cortical travelling waves, in both forwards/backwards and lateral directions. Our findings showed that agreeableness is related to the strength of interhemispheric alpha waves travelling from electrodes over the right temporal regions, and that this relation was largely attributable to the compassion (rather than politeness) aspect of agreeableness. They also showed that openness/intellect was related to the strength of backwards travelling alpha waves down midline electrodes, and that this relation was significant for openness but not intellect. These relationships were relatively strong compared to those commonly reported in the personality neuroscience literature using sufficiently powered sample sizes, with strong Bayesian evidence in favour of the relationship between openness/intellect and backwards travelling waves, and decisive Bayesian evidence in favour of the presence of the relationships to agreeableness / compassion (Jeffreys, 1998). Furthermore, we assessed the potential replicability of our findings by randomly excluding 25 or 50% of our sample and re-testing the correlations. These validation analyses were consistent in showing a relationship between agreeableness and rightwards travelling waves, and 19/20 of these random sub-sample tests showed a significant effect for the relationship between openness/intellect and backwards travelling waves. This demonstrates that the effects were not dependent on a specific selection of participants for the study, indicating that our findings would likely replicate in an independent sample. The strength and novelty of our results provides important new insights into potential neural mechanisms underlying personality traits.

Our findings build on previous research that has shown links between personality traits and neural oscillations (Pacheco et al., 2024). However, that previous research used methods that do not indicate whether the oscillations might be travelling. Although links between personality traits and standing neural oscillations are in line with the view that personality differences must be underpinned by stable differences in neural activity (Allen & DeYoung, 2017), they shed little light on the mechanistic reasons for these neural correlates of personality traits. Our current findings extend this previous research to demonstrate relationships between specific personality traits and the strength of alpha travelling waves in specific directions. In particular, individuals who scored higher in openness/intellect (particularly in the openness aspect) showed stronger backwards travelling waves from frontal to posterior midline electrodes, while individuals who scored higher in agreeableness (particularly in the compassion aspect) showed stronger rightwards travelling interhemispheric waves along the central electrode line. These findings align with other research that has suggested measures of connectivity between brain regions might be particularly informative in personality neuroscience, and our results show broad consistency with fMRI connectivity patterns reported in previous research (Adelstein et al., 2011).

The travelling wave patterns we assessed have been shown to provide an index of information flow through the cortex (Bailey, Hill, Godfrey, Perera, Hohwy, et al., 2024; Lozano-Soldevilla & VanRullen, 2019; Mohan et al., 2024; Muller et al., 2014; Pang et al., 2020). Specifically, travelling waves have been interpreted as reflecting the propagation of prediction errors up the cortical hierarchy, and the allocation of top-down prediction weighting from higher levels in the cortical hierarchy to lower levels in the cortical hierarchy (Alamia et al., 2023). Alpha cortical travelling waves have also been shown to inhibit gamma activity and neuronal spiking, with the inhibition functioning to supress predicted signals across distant cortical regions, perhaps explaining the mechanism by which travelling waves fulfil predictive processing functions (Xiong et al., 2024). Macro-scale alpha travelling waves (as detected in our study) have also been shown to modulate micro-scale travelling waves, which in turn have been shown to synchronise neuronal spiking (Sreekumar et al., 2020). This suggests that travelling waves provide a role in synchronising information processing across the cortex, with this regulatory effect on processing perhaps providing a putative mechanism by which travelling waves underpin variations in personality traits. Our findings therefore suggest that the relative strength of information flow between certain regions of the cortex is related to specific personality traits. Interpreting the location and direction of the cortical travelling wave patterns that relate to personality traits through this lens, coupled with prior evidence for the specialisation of specific brain regions, may therefore offer unique insight into the neural sources of individual differences in personality.

Through this lens, it is intriguing that Agreeableness (especially compassion) is related to cortical travelling waves that propagate from the right hemisphere to the left hemisphere, with an origin detected in electrodes over temporal and temporoparietal brain regions. Agreeableness is characterised by altruism, empathy, and cooperation, and is associated with social cognition and perspective taking (Graziano et al., 2007; Nettle & Liddle, 2008). Research indicates that the superior temporal region is associated with interpreting the actions and intentions of others based on their movements (Pelphrey & Morris, 2006). In alignment with this, research using functional magnetic resonance imaging has shown that agreeableness correlates with volumes in the left superior temporal region (Li et al., 2017, cf, Allen et al., 2017). Activity in the right temporoparietal junction has been shown to be associated with the processing of empathy, sympathy, taking another’s perspective, and processing social cues (Decety & Lamm, 2007). Research using fMRI also indicates that connectivity to the right temporoparietal junction was predictive of agreeableness for the majority of individuals (Nostro et al., 2018). Furthermore, myelin-based microstructural connectivity in an interhemispheric network that involved the right temporoparietal junction has been shown to both relate to agreeableness and mediate the better life satisfaction associated with agreeableness (Wu et al., 2024). Our data did not enable accurate source analysis, prohibiting specific localisation of the rightwards travelling wave origins. Thus, given the effects of volume conduction on the EEG signal, we are unable to draw conclusions about the source regions of the cortical travelling waves. Nevertheless, the consistency between the functional associations reported across these general brains and our findings that cortical travelling waves propagate more strongly from electrodes lying over these regions in participants who score highly in trait compassion provides an intuitive link between established neuroanatomic specialisation and a personality trait. This consistency between the information flow direction suggested by our findings and knowledge of brain region function suggests a potential mechanistic pathway between neural activity, behaviour, and personality traits.

The pathway by which this pattern originates is likely to be driven by both genetic and environmental influences. Potential pathways could involve: 1) individuals who inherit genes that influence their brain activity towards stronger propagation of activity from certain brain regions may be more likely to be motivated by consideration of others, resulting in an increased frequency or intensity of thoughts, emotions and behaviours related to compassion, leading to their endorsement of more compassionate items on the personality questionnaire; 2) individuals who experience life events that cause them to be motivated by compassionate concern for others develop more synaptic connections from these brain regions, leading to stronger cortical travelling waves propagating from those regions while at rest, providing a readiness for those connections to be activated, perhaps driving compassionate behaviour in the future, and leading to higher endorsement of higher scores on compassion questions when answering a personality questionnaire; or 3) most likely, a combination of both 1 and 2 (with different degrees of influence within different individuals). These pathways may then have led to an increased probability of rightwards travelling waves during the resting state, due to increased proclivity of individuals high in agreeableness to rest in a state that involves engaging these travelling waves and relevant networks, in alignment with suggestions that personality traits correspond to probability distributions of mental states (Fleeson & Gallagher, 2009). Alternatively (or additionally), the differences may be enduring, and may influence behaviour on a daily basis, in alignment with more biologically oriented theories of personality (e.g. Depue & Morrone-Strupinsky, 2005). In support of this, travelling wave directions have been shown to structural connectivity gradients in the human connectome detected by MRI (Koller et al., 2024), suggesting the travelling wave patterns we detected may reflect traits that persist outside of the resting period (rather than being state dependent).

Turning now to openness/intellect, individuals higher on this trait tend to be characterised by abstract thinking, cognitive flexibility, and curiosity (DeYoung et al., 2014; Kaufman et al., 2016). Interestingly, our analyses indicated that backwards travelling waves were reliably associated with openness, but not intellect. While openness describes engagement in perception, fantasy, aesthetics, and emotion, intellect reflects engagement with abstract and semantic information via reasoning (DeYoung, 2015). Backwards travelling waves have previously been reported to indicate the voluntary control of attention (Alamia & VanRullen, 2019; Pang et al., 2020), which falls under the umbrella of executive functions, and has been shown to relate to the intellect aspect of openness/intellect rather than the openness aspect (Smillie et al., 2016). However, research into mindfulness meditators has suggested that while participants are resting during EEG recordings, backwards travelling waves might reflect thoughts about the future or past, or daydreaming, rather than the executive functions reflected by backwards waves during a cognitive task (Bailey, Hill, Godfrey, Perera, Hohwy, et al., 2024). In this context, the stronger backwards travelling wave strength in individuals who scored higher in trait openness might reflect the engagement of daydreaming during the resting EEG recording, eliciting an increase in backwards waves, reflective of top-down processes that are not necessarily indicative of the executive functions that might be associated with the intellect aspect of openness/intellect. This pattern of results aligns with fMRI research indicating that openness relates to the efficiency of activation of the default mode network (Beaty et al., 2016), as well as connectivity to core midline hubs of the default mode network (Adelstein et al., 2011)—brain regions that are associated with cognitive flexibility, creativity, and imagination.

While the implications of our results are predominantly focused on improving our understanding of the neural sources of personality traits, we note our results have potential practical applications in the area of personality change interventions. According to some estimates, around two thirds of people would like to change some aspect of their personalities—a finding that is reflected in the prominence of the self-improvement industry (Miller et al., 2019). There is also growing evidence that personality traits can be changed through intervention (Bleidorn et al., 2019). Recent proof-of-concept research has suggested that cortical travelling waves can be modulated by a modified transcranial alternating current stimulation device with two stimulation points offset by a brief delay (Lee et al., 2023). Future research might therefore explore brain stimulation methods targeting cortical travelling waves as a potential avenue for increasing trait levels of compassion or openness, for individuals seeking to do so.

## Limitations

Although our findings indicated decisive Bayesian evidence for a relationship between agreeableness/compassion and rightwards travelling waves, and strong evidence for a relationship between openness and backwards travelling waves, the strength of our conclusions for this effect are limited by a few considerations. First, the current study was exploratory in nature, using a data driven approach, rather than hypothesis driven. Such a data driven approach is in one sense a strength, enabling the detection of relationships that are unlikely to be conceived of via purely hypothesis driven approaches (Bailey, Fulcher, et al., 2024). Additionally, despite our testing of many correlations between personality traits and different travelling wave directions, these tests were controlled for false positive inflation by multiple comparisons both within relationships to each personality factor using cluster statistics, and across the different personality factors using the false discovery rate control. Our results were also robust to different random splits of the data into subsamples, suggesting a real effect that would likely replicate. However, the hypothesis driven testing of our results on an independent dataset would provide full confidence that our results reflect a true finding. This is particularly important given that the data we used in this analysis has provided results after analysis of non-travelling oscillations that have not replicated in an independent sample (Fruehlinger et al., 2024).

Second, it may be that the EEG recording session itself acted as a conditioning stimulus, prompting certain thoughts, emotions, or states of mind, and that these factors may have interacted with personality traits to lead to the findings we have reported. If this is the case, our findings might not reflect stable processes underlying agreeableness or openness, but rather a proclivity towards specific mental states that are associated with specific neural travelling wave markers and elicited only within specific contexts—in this case, a neuroscientific study (Jach et al., 2020). Even if this were the case, it would not negate the finding that compassion and openness were linked with travelling waves. Given the myriad of potential interacting influences on participants in the paradigm, no specific influence seems more likely than the interpretation that our results reflect a neural marker of the two personality traits (Jach et al., 2020). Additionally, travelling wave directions have been shown to follow structural connectivity gradients (Koller et al., 2024), suggesting the travelling wave patterns we detected may reflect traits that persist outside of the resting period. Future research might explore whether stronger travelling waves observed at rest are related to stronger travelling waves during tasks, which might indicate that resting travelling wave strengths are trait markers that have an influence on behaviour.

Third, previous research on the same data has indicated that agreeableness can be decoded from non-oscillatory activity (which shows a 1/f log-power log-frequency distribution) after excluding oscillatory activity (Pacheco et al., 2024). Additionally, the same research indicated that measures of standing oscillatory activity alone (after removing the influence of non-oscillatory 1/f activity) provided better decoding of neural activity related to personality traits than measures of standing broadband activity (Pacheco et al., 2024). This finding of increased sensitivity after separating oscillatory and non-oscillatory EEG activity is common across different populations (e.g. McQueen et al., 2024), highlighting the potential importance of addressing the potential confound of 1/f non-oscillatory activity. We measured cortical travelling waves using 2D and 3D-FFTs. While these techniques allowed the novel measure of cortical travelling waves in both forwards/backwards and lateral directions across the scalp, FFT analyses (including 2D-FFT and 3D-FFT analyses) do not separate oscillatory activity from non-oscillatory activity. As such, our results do not indicate whether the patterns we observed are driven by travelling waves that do indeed oscillate (temporally), or simply voltage shifts within the alpha temporal frequency that travel across the scalp. It may be possible for future research to address this using travelling wave analyses methods that control for the influence of non-oscillatory activity, but we are not aware of such methods currently. Despite noting this limitation, it is worth mentioning that although temporal oscillatory measures might be confounded by non-oscillatory 1/f activity, there is currently no indication that travelling waves are vulnerable to the same confound. Additionally, we focused on the alpha frequency. Alpha oscillations show the largest peak above non-oscillatory 1/f activity, and previous research has shown that when analyses were restricted to alpha peaks above the non-oscillatory 1/f activity, alpha travelling waves were still prominent (Zhang et al., 2018). Furthermore, recent modelling and empirical work has suggested that unless neurophysiological effects involve differences in the decay time of synaptic currents, correcting for the non-oscillatory 1/f activity can introduce measurement errors that are larger than the potential gains in the measurement accuracy of oscillations that can be obtained by correcting for the non-oscillatory 1/f activity (Brake et al., 2024).

Finally, the data used in the current analysis was recruited from a WEIRD population (white, educated, industrialized, rich and democratic) (Pacheco et al., 2024). As such, it is not clear that our results would generalise to other populations. Future research in other populations is required to determine whether the relationships between personality traits and cortical travelling waves in different cultures, in alignment with calls for the future of EEG research to be more inclusive and representative (Mushtaq et al., 2024).

## Acknowledgements

We are grateful to Hayley Jach, Tianru Sun, Vu Ngoc Duong, and Elizabeth Robinson for assistance with data collection. We thank Olivia Carter for helpful feedback on a draft of this manuscript.

