## Supplementary Materials for "Cortical travelling waves may underpin variation in personality traits"

**Eyes-closed data showed the same relationships as eyes-open data**

In addition to the relationship between right to left travelling waves and Agreeableness in the eyes open data, we tested for a relationship in the eyes closed resting data to determine the consistency of the effect across different states. The correlation between Agreeableness and eyes closed resting right to left traveling waves at the highest spatial frequency and 9-10Hz was significant (rho = 0.236, p < 0.001, BF10 = 7.467). The relationship was also significant for compassion (rho = 0.246, p < 0.001, BF10 = 14.862) and for politeness, although with Bayesian evidence against the effect (rho = 0.147, p = 0.011, BF10 = 0.324). A linear regression including compassion and politeness as predictors for the rightwards cortical travelling wave strengths in the eyes-closed resting data was also undertaken. This analysis showed a significant effect for compassion (t = 2.841, p = 0.005) but not for politeness (t = 0.535, p = 0.593), suggesting the effect was specific to compassion, similar to our analysis of the eyes open resting variables. This result indicates that the patterns are consistent across different resting states.

Similarly, in addition to the relationship between backwards travelling waves and Openness/Intellect in the eyes open data, we tested for a relationship in the eyes closed resting data. The correlation between Openness/Intellect and eyes closed resting backwards traveling midline waves at the lowest spatial frequency and 10-11Hz was significant, although with only weak Bayesian evidence in support of the relationship (rho = 0.157, p = 0.007, BF10 = 1.766). The relationship was also significant for openness (rho = 0.196, p < 0.001, BF10 = 23.096), but not for intellect, with Bayesian evidence against the effect (rho = 0.055, p = 0.349, BF10 = 0.096). This result indicates that the patterns were consistent across different resting states.

**Analysis of latent personality variables showed the same patterns as our primary analyses**

Next, to assess the robustness of our results to different analysis methods, we tested relationships between the cortical travelling waves that showed significant relationships to personality traits in our primary analyses to latent personality factors. The same pattern of associations between rightwards travelling waves and Agreeableness (and its aspects) was evident when personality traits were estimated as latent variables: Waves travelling from the right to the left along the C line were significantly correlated with latent Agreeableness (rho = 0.283, p < 0.001, BF10 = 1317.217). A similar correlation was present with latent compassion (rho = 0.270, p < 0.001, BF10 = 436.566), which was stronger than the effect for latent politeness (rho = 0.183, p = 0.002, BF10 = 2.112). A linear regression including latent compassion and latent politeness as predictors for the rightwards cortical travelling wave strengths was also undertaken. This analysis showed a significant effect for compassion (t = 3.466, p < 0.001) but not for politeness (t = 1.036, p = 0.301), suggesting the effect was specific to latent compassion, similar to our analysis of the non-latent variables.

The patterns noted for Openness/Intellect were also similar for the latent variables, but weaker: latent Openness/Intellect was positively correlated with backwards travelling wave strength (rho = 0.137, p = 0.018). The same correlation was present for the latent variable of Openness (rho = 0.171, p = 0.003), but again, not for the latent variable of Intellect (rho = 0.088, p = 0.131); the difference between these correlations was statistically non-significant, 𝑧 = 1.186, p = 0.236. A linear regression including these two variables as predictors for the backwards cortical travelling wave strengths confirmed a significant effect only for openness (t = 2.715, p = 0.007) but not for intellect (t = 0.954, p = 0.341).

**The centroparietal and frontocentral electrode lines also showed similar relationships to agreeableness**

The relationship between Agreeableness and rightwards travelling waves was also significant when our 2D-FFT was computed using the fronto-central, and centroparietal lateral electrode lines. Waves travelling from the right to the left at the maximum spatial frequency along the centroparietal electrode line were significantly correlated with Agreeableness (rho = 0.214, p < 0.001, BF10 = 21.592). A similar correlation was present with compassion (rho = 0.224, p < 0.001, BF10 = 42.512), and with politeness (although weaker, with Bayesian evidence against the relationship: rho = 0.134, p = 0.021, BF10 = 0.502). A linear regression including compassion and politeness as predictors for the rightwards cortical travelling wave strengths was also undertaken. This analysis showed a significant effect for compassion (t = 3.083, p = 0.002) but not for politeness (t = 0.671, p = 0.503), suggesting the effect was specific to compassion, similar to our analysis of the non-latent variables.

The correlations were also significant when testing relationships in the fronto-central electrode line, although of weaker strength and with lower Bayesian evidence than for the central and centroparietal lines. Waves travelling from the right to the left along the FC line were significantly correlated with Agreeableness (rho = 0.204, p < 0.001, BF10 = 3.537). A similar correlation was present with compassion (rho = 0.204, p = 0.011, BF10 = 1.864), and with politeness (although weaker: rho = 0.140, p = 0.016, BF10 = 0.608). A linear regression including latent compassion and latent politeness as predictors for the rightwards cortical travelling wave strengths was also undertaken. This analysis showed a non-significant trend towards a significant effect for compassion (t = 1.919, p = 0.056) but not for politeness (t = 1.189, p = 0.235).

**Using the mean instead of the 75th percentile showed the same relationships**

Additionally, despite our rationale that testing the 75th percentile of travelling wave strengths across epochs would be more likely to enable detection of relationships to personality traits due to these epochs representing stronger engagement of neural processes of interest, it is more typical to test the mean across epochs. To assess the consistency of our results when applying more typical travelling wave quantification methods, we repeated tests of relationships between personality traits and our 2D-FFT outputs using mean values across epochs. These analyses showed the same pattern of results and significant effects as our primary analyses. The correlation between Agreeableness and the mean eyes-open resting right to left traveling waves at the highest spatial frequency and 9-10Hz was significant (rho = 0.205, p < 0.001, BF10 = 15.433). The relationship was also significant for compassion (rho = 0.243, p < 0.001, BF10 = 141.583), but not for politeness, with Bayesian evidence against the effect (rho = 0.103, p = 0.076, BF10 = 0.208). A linear regression including compassion and politeness as predictors for the mean rightwards cortical travelling wave strengths in the eyes-open resting data was also undertaken. This analysis showed a significant effect for compassion (t = 3.659, p < 0.001) but not for politeness (t = -0.030, p = 0.976), suggesting the effect was specific to compassion, similar to our analysis of the effects obtained by testing the values at the 75th percentile.

Furthermore, the correlation between openness/intellect and mean eyes-open resting backwards traveling waves at the lowest spatial frequency and 10-11Hz was significant (rho = 0.182, p = 0.002, BF10 = 5.794). The relationship was also significant for openness (rho = 0.220, p < 0.001, BF10 = 18.120), but not for intellect, with Bayesian evidence against the effect (rho = 0.079, p = 0.178, BF10 = 0.207). A linear regression including openness and intellect as predictors for the backwards cortical travelling wave strengths in the eyes-open resting data was also undertaken. This analysis showed a significant effect for openness (t = 3.090, p = 0.002) but not for intellect (t = 0.652, p = 0.515), suggesting the effect was specific to openness, similar to our analysis of the effects obtained by testing the values at the 75th percentile.

**Results for agreeableness were robust against different normalisation approaches**

Despite these interesting results, inspection of the data revealed that the decibel values for the rightwards lateral travelling waves at the highest spatial frequency were typically negative after division by the mean of 100 null shuffles. This suggests that the null shuffles of electrode orders contained more power than the real data in these highest lateral spatial frequencies, requiring consideration of whether the positive correlations we detected were driven by the presence of true lateral travelling waves. To assess whether our result in these highest lateral spatial frequency cells may have been driven by the normalisation procedure (which would suggest an effect driven by non-spatial patterns), we tested correlations between agreeableness, compassion, and the mean null shuffle versions of the data. If these correlations were significant, it would demonstrate that the normalisation against the null shuffles was driving the significant correlations we have reported for rightward travelling waves. The correlation between the null shuffle versions of the data and agreeableness was not significant (rho = 0.043, p = 0.463, BF10 = 0.110), and nor was the relationship to compassion (rho = 0.017, p = 0.775, BF10 = 0.077). To further assess whether our result in these highest lateral spatial frequency cells may have been influenced by the normalisation procedure, we tested correlations against agreeableness on the log10 transformed versions of the real data, after normalisation against stationary (1D-FFT) mean alpha power across the electrodes of interest (instead of the null shuffled versions of the electrode order) (Zeng et al., 2024). This provided control for variations in standing wave alpha power (in the temporal but not spatial domain), while still measuring the cortical travelling wave strength (Zeng et al., 2024). Within these tests, the correlation between agreeableness and power in the highest right to left travelling spatial frequency from 9-10Hz was still significant, (rho = 0.262, p < 0.001, BF10 = 611.413), as was the correlation with compassion (rho = 0.256, p < 0.001, BF10 = 150.596).

To further address this potential issue, we also tested for correlations after normalising data to null shuffles obtained by randomly swapping electrodes between epochs. This approach destroyed the spatiotemporal relationships that produced travelling waves, as the timing of oscillations between epochs would not be synchronised, but preserved both variations in alpha power, and variations in the relationship between electrodes in alpha power. These correlations were also significant - the correlation between agreeableness and power in the highest right to left travelling spatial frequency from 9-10Hz was still significant, (rho = 0.205, p < 0.001, BF10 = 48.979), as was the correlation with compassion (rho = 0.204, p < 0.001, BF10 = 21.371). Interestingly, normalising using this method demonstrated that the rightward travelling waves did exceed the null shuffled data in many participants, but not all participants, with a mean value of 0.170 dB (SD = 0.306, minimum = -0.786, maximum = 0.885). This suggests that on average, the real rightwards travelling wave strength exceeded the strength of the null shuffles, but not by much, and not for every participant. This pattern indicates that although the rightwards travelling wave does reflect a true signal in the data, the signal is weak and not present in all individuals. As such, these tests confirm that the effects were indeed driven by a relationship between true rightwards travelling wave strength. True rightwards travelling waves were present more commonly in individuals scoring higher in agreeableness, and individuals higher in agreeableness showed rightwards travelling waves that exceeded the values obtained via null shuffles of the data that destroyed the travelling wave patterns by a larger amount compared to individuals scoring lower in agreeableness and compassion.

Additionally, previous research on a subset of the dataset used in the current study has shown that agreeableness is negatively correlated with posterior alpha power (Jach et al., 2020). As such, we performed an additional test to assess the potential that our results might be simply driven by differences in alpha power. In this test, we z-score transformed each electrode’s time series separately prior to the 2D-FFT. This normalises for differences in amplitude between electrodes and between individuals, controlling for potential differences in alpha power, while preserving the spatial properties of the cortical travelling waves (since the phase angles between the electrodes are preserved by this transform). After these computations, the correlation between Agreeableness and travelling waves from the far right at 9-10Hz was still significant, and in fact even stronger than the initial tests (rho = 0.274, p < 0.001, BF10 = 741.186). The correlation was also significant for compassion (rho = 0.295, p < 0.001, BF10 = 1268.632), and for politeness, although again weaker than for compassion (rho = 0.175, p = 0.003, BF10 = 2.302). This indicates that the relationships were present even after controlling for potential differences in alpha power across the electrodes.

**Analyses of subsets of lateral electrodes indicates results for agreeableness were driven by interhemispheric travelling waves**

Finally, we note that the analyses of lateral travelling waves reported thus far do not reveal whether the waves travel within each hemisphere, or whether they travel between hemispheres. To address this, we performed additional 2D-FFTs that 1) only included the right hemisphere central electrodes (T8 to Cz), 2) included midline electrodes only (C3 to C4), 3) included lateral electrodes, but not temporal electrodes (C5 to C6), and 4) included all central line electrodes (T7 to T8, 9 electrodes). Relationships between Agreeableness and lateral cortical travelling waves restricted to the right hemisphere only were not significant at any spatial frequency (all p > 0.10 when only electrodes from T8 to Cz were included in the 2D-FFT). This suggests that our primary results were not driven by waves travelling from the right-most electrodes to the midline. Similarly, when our 2D-FFT was restricted to electrodes from C3 to C4, no significant correlations were present at any spatial frequency (all p > 0.10). Only when we included electrodes from C5 to C6 did significant effects become apparent, although the effects within this analysis were weaker than our results from analyses that included T7 and T8 electrodes. The analysis including electrodes from C5 to C6 showed a significant correlation between Agreeableness and right to left travelling waves at the middle spatial frequency and 9-10Hz (rho = 0.165, p = 0.004, BF10 = 3.824). The analysis that included the 9 central line electrodes (T7 to T8) showed a significant correlation between agreeableness and right to left travelling waves at the second lowest spatial frequency (rho = 0.269, p < 0.001, BF10 = 540.882). Given that the list of electrodes from T8 to Cz and from C3 to C4 both contained five electrodes (the same number of lateral electrodes as in the 3D-FFT and our post-hoc 2D-FFTs), the lack of effect when testing the right hemisphere alone and midline electrodes is unlikely to be due to a reduced number of electrodes. Therefore, the pattern of results suggests that the relationship between right to left cortical travelling waves and agreeableness / compassion is produced by interhemispheric cortical travelling wave patterns rather than travelling waves from lateral electrodes to midline electrodes.
